# When do longer reads matter? A benchmark of long read de novo assembly tools for eukaryotic genomes

**DOI:** 10.1101/2023.01.30.526229

**Authors:** Bianca-Maria Cosma, Ramin Shirali Hossein Zade, Erin Noel Jordan, Paul van Lent, Chengyao Peng, Stephanie Pillay, Thomas Abeel

## Abstract

**Background:** Assembly algorithm choice should be a deliberate, well-justified decision when researchers create genome assemblies for eukaryotic organisms from third-generation sequencing technologies. While third-generation sequencing by Oxford Nanopore Technologies (ONT) and Pacific Biosciences (PacBio) have overcome the disadvantages of short read lengths specific to next-generation sequencing (NGS), third-generation sequencers are known to produce more error-prone reads, thereby generating a new set of challenges for assembly algorithms and pipelines. Since the introduction of third-generation sequencing technologies, many tools have been developed that aim to take advantage of the longer reads, and researchers need to choose the correct assembler for their projects.

**Results:** We benchmarked state-of-the-art long-read *de novo* assemblers, to help readers make a balanced choice for the assembly of eukaryotes. To this end, we used 13 real and 72 simulated datasets from different eukaryotic genomes, with different read length distributions, imitating PacBio CLR, PacBio HiFi, and ONT sequencing to evaluate the assemblers. We include five commonly used long read assemblers in our benchmark: Canu, Flye, Miniasm, Raven and Redbean. Evaluation categories address the following metrics: reference-based metrics, assembly statistics, misassembly count, BUSCO completeness, runtime, and RAM usage. Additionally, we investigated the effect of increased read length on the quality of the assemblies, and report that read length can, but does not always, positively impact assembly quality.

**Conclusions:** Our benchmark concludes that there is no assembler that performs the best in all the evaluation categories. However, our results shows that overall Flye is the best-performing assembler, both on real and simulated data. Next, the benchmarking using longer reads shows that the increased read length improves assembly quality, but the extent to which that can be achieved depends on the size and complexity of the reference genome.

## Introduction

*De novo* genome assembly is essential in several leading fields of research, including disease identification, gene identification, and evolutionary biology [1–4]. Unlike reference-based assembly, which relies on the use of a reference genome, de novo assembly only uses the genomic information contained within the sequenced reads. Since it is not constrained to the use of a reference, high quality de novo assembly is essential for studying novel organisms, as well as for the discovery of overlooked genomic features, such as gene duplication [5], in previously assembled genomes.

The introduction of Third Generation Sequencing (TGS) led to massive improvements in *de novo* assembly. The advent of TGS has addressed the main drawback of Next Generation Sequencing (NGS) platforms, namely the short read length, but has introduced new challenges in genome assembly, because of the higher error rates of long reads. The leading platforms in long-read sequencing are Pacific Biosciences Single Molecule, Real-Time sequencing (often abbreviated as “PacBio”) and Oxford Nanopore (ONT) sequencing [6].

Since the introduction of TGS platforms, many methods have been developed that aim to take the most benefits from the longer read length and overcome the new challenges due to sequencing error. Recent studies have been conducted to compare long-read de novo assemblers. One such study was conducted by Wick and Holt [7], who focused on long-read de novo assembly of prokaryotic genomes. Eight assemblers were tested on real and simulated reads from PacBio and ONT sequencing, and evaluation metrics included sequence identities, circularisation of contigs, computational resources, as well as accuracy. Murigneux et al. [8] performed similar experiments on the genome of *M. jansenii*, although in this case, the focus was on comparatively benchmarking Illumina sequencing and three long-read sequencing technologies, in addition to the comparison of long-read assembly tools. Studies narrowed down to just one type of sequencing technology include those of Jung et al. [9], who evaluated assemblers on real PacBio reads from five plant genomes, and Chen et al. [10], who used Oxford Nanopore real and simulated reads from bacterial pathogens in their comparison. Except for the Wick and Holt study, which provides a compressive comparison on de novo assembly of prokaryotic genomes, other studies are either comparing the assemblers on single genome or using data from a single sequencing platform. Here, we provide a comprehensive comparison on de novo assembly tools on all TGS technologies and 7 different eukaryotic genomes, to complement the study of Wick and Holt.

In this study, we are benchmarking these methods using 13 real and 72 simulated datasets (see Figure 1) from both PacBio and ONT platforms to guide researchers to choose the proper assembler for their studies. Benchmarking using simulated reads allows us to accurately compare the final assembly with the ground truth, and benchmarking using the real reads can validate the results based on simulated reads. The assembler comparison presented in this manuscript complements the literature that has already been published, by introducing an analysis of not just assembler performance, but also of the effect of read length on assembly quality. Although increased read length is considered an advantage, we investigate if it is always a necessary advantage to have for assembly performance. To that end, the scope of the study extends to six model eukaryotes that provide a performance indication for genomes of variable complexity, covering a wide range of taxa on the eukaryotic branch of the Tree of Life [11]. Complexity in genome assembly is determined by multiple variables, the most notable of which is the proportion of repetitive sequences within the genome of a particular organism. Complexity in eukaryotic genomes is further exacerbated by size and organization of chromosomal architecture, including telomeres and centromeres, and the presence of circular elements such as mitochondrial and chloroplast DNA.

**Figure 1:**
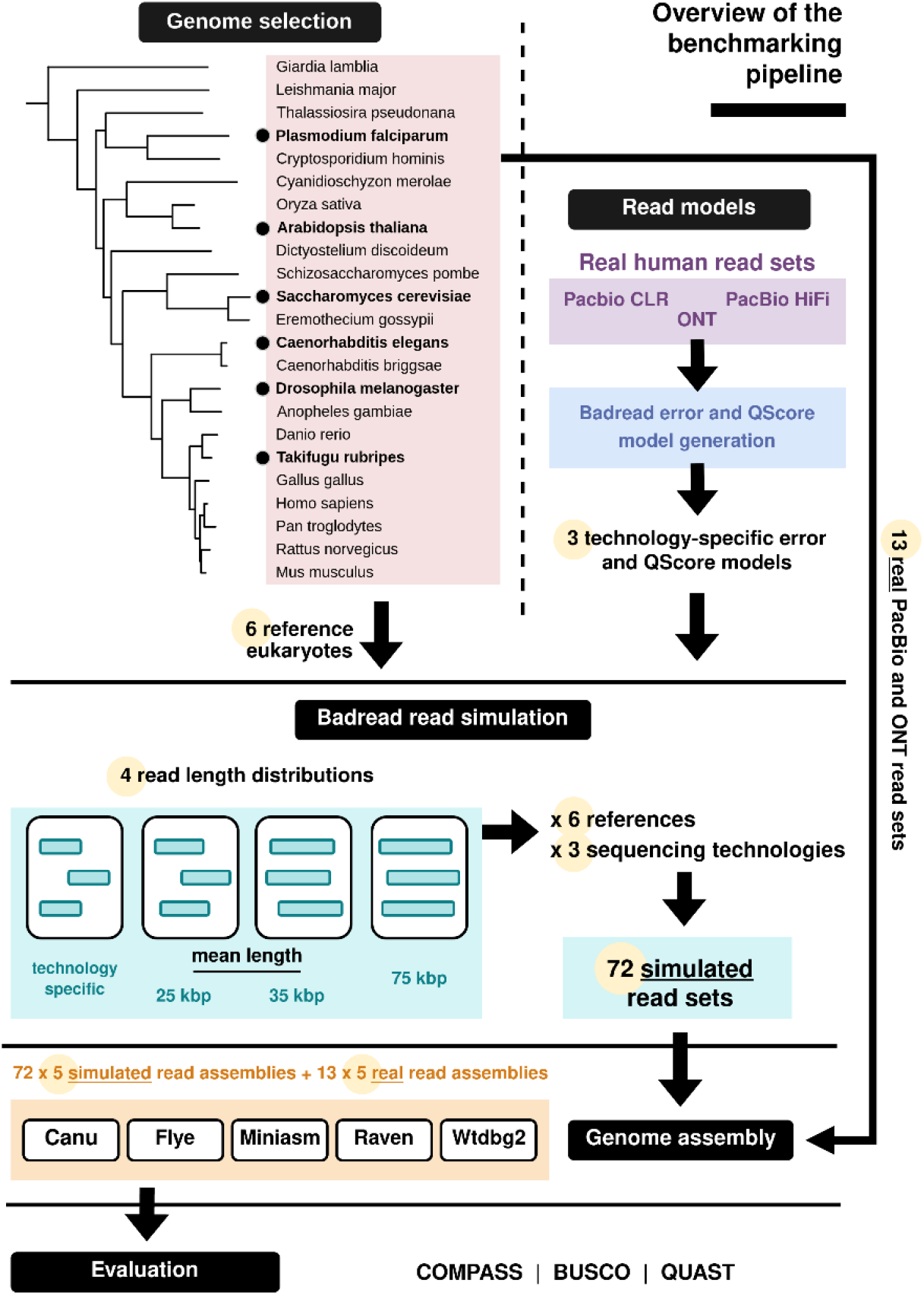
The benchmarking pipeline. We first select 6 representative eukaryotes from the Tree of Life (Letunic and Bork, 2021) and use Badread’s error and QScore model generation feature (Wick, 2019) to create 3 models of state-of-the-art long sequencing technologies. This is input to the read simulation stage, where we simulate reads from all genomes, with four different read length distributions. We then perform assembly of simulated and real reads, using five long-read assemblers. Lastly, we evaluate all assemblies based on several criteria.

De novo genome assembly evaluation remains challenging, as it represents a process that must account for variables such as the goal of an assembly and the existence of a ground-truth reference. A standard evaluation procedure was introduced in the literature by the two Assemblathon competitions [12,13], which outlined a selection of metrics that encompasses the most relevant aspects of genome assembly, however, these metrics require a reference sequence. Most of these metrics are adopted in our benchmark.

Consequently, this study addresses two main objectives. First, we provide a systematic comparison of five state-of-the-art long-read assembly tools, documenting their performance in assembling real and simulated PacBio Continuous Long Reads (CLRs), PacBio Circular Consensus Sequencing (CCS) HiFi reads, and Oxford Nanopore reads, generated from the genomes of *S. cerevisiae, P. falciparum, C. elegans, A. thaliana, D. melanogaster*, and *T. rubripes*. Our second objective is to investigate whether increased read length has a positive effect on overall assembly quality, given that increasing the length of reads is an on-going effort in the development of Third Generation Sequencing platforms [14].

## Materials and methods

### Data

In this study, we are using real and simulated data from various organisms to benchmark long read *de novo* assembly tools.

#### Reference genomes

We selected six reference genomes from eukaryotic organisms represented in the Interactive Tree Of Life (iTOL) v6 [11]: *S. cerevisiae* (strain S288C), *P. falciparum* (isolate 3D7), *C. elegans* (strain VC2010), *A. thaliana* (ecotype Col-0), *D. melanogaster* (strain ISO-1), and *T. rubripes*. Assembly accessions are included in Supplementary Table S1.

The reference assemblies for *C. elegans, D. melanogaster*, and *T. rubripes* included uncalled bases. In these cases, before read simulation, each base N was replaced with base A, as done by Wick and Holt [7]. This avoids ambiguity in the read simulation process and consequently simplifies the evaluation of the simulated-read assemblies. As such, we used this modified version as a reference when evaluating all assemblies of simulated reads from these four genomes. In the evaluation of real-read assemblies, the original assemblies were used as references.

#### Simulated reads

All simulated read sets were generated using Badread v0.2.0 [15]. To create read error and QScore (quality score) models in addition to the simulator’s own default models, Badread requires the following three parameters: a set of real reads, a high-quality reference genome, and an alignment file, obtained by aligning the reads to the reference genome. We used real read sets from the human genome to create error and QScore models that reflect the state-of-the-art for three sequencing technologies: PacBio Continuous Long Reads (CLRs), PacBio Circular Consensus Sequencing (CCS) HiFi reads, and Oxford Nanopore reads.

To create the models, we used the real read sets sequenced from the human genome and aligned to the latest high-quality human genome reference assembled by [16]: assembly T2T-CHM13v2.0, with RefSeq accession GCF_009914755.1. The alignment was performed using Minimap2 v2.24 [17] with default parameters. The sources for these sequencing data are outlined in Supplementary Table S2, as well as the read identities for each technology, which are later passed as parameters for the simulation stage.

For each of the six reference genomes, we simulated reads that imitate PacBio CLR, PacBio HiFi, and Oxford Nanopore sequencing, with four different read length distributions, using Badread. The first read simulation represents the current state of the three long-read technologies. The other three simulations reflect data points in-between technology-specific values and ultra-long reads, data points of a similar length as ultra-long-reads, and longer than ultra-long reads. Since Badread’s read length models are parameterized by gamma distributions, we need to define the mean and standard deviation of the gamma distributions for these simulations. The values for the mean and standard deviation of these distributions were selected as follows. First, we calculated the read length distributions of the real read sets in Supplementary Table S2 and simulated an initial iteration of reads using these technology-specific values. For choosing these values for the other three iterations, we analysed a set of Oxford Nanopore Ultra-Long reads used in the latest assembly of the human genome (Nurk *et al*., 2022). We selected GridION run SRR12564452, available as sequence data in BioProject PRJNA559484, with a mean read length of approximately 35.7 kbp, and a standard deviation of 42.5 kbp.

A full overview of the mean and standard deviation of all four read length distributions is given in Table 1. Note that, for each of the technologies, the standard deviation for the last three distributions was derived from the mean, using the ratio between the mean and standard deviation reflected by the technology-specific values. Hence, for the last three iterations, the mean read length is consistent across sequencing technologies, but the standard deviation varies.

**Table 1:**
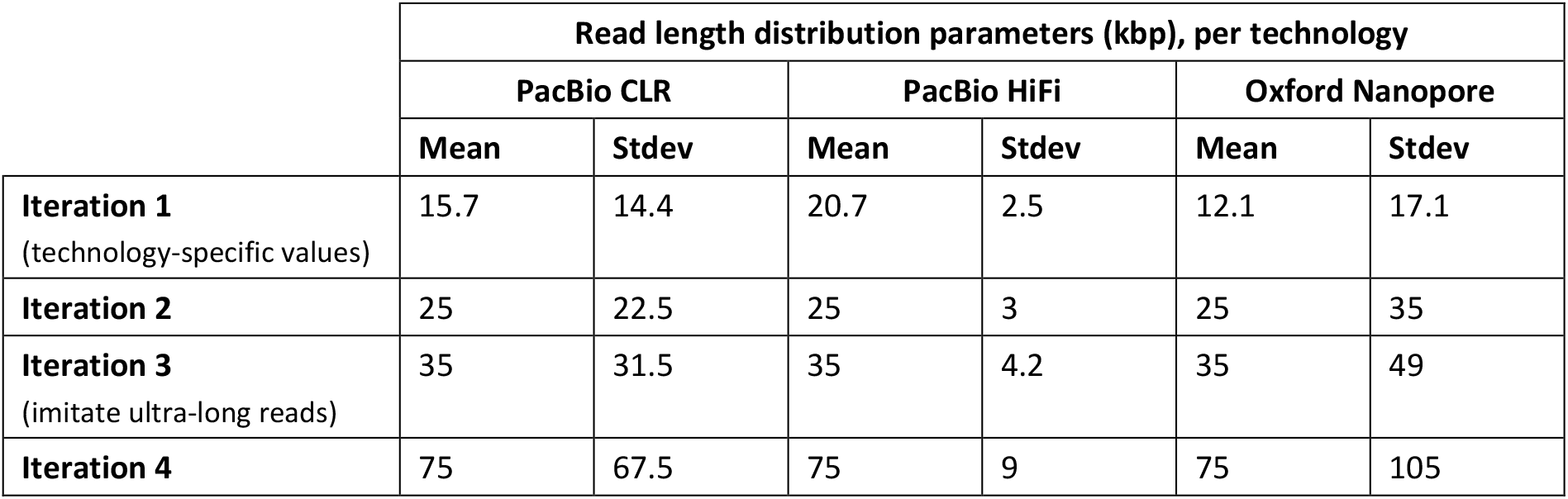
The mean and standard deviation describing the read length distributions used in our simulations. Note that read length increases with each iteration, and the distribution parameters are different for each technology.

|  | Read length distribution parameters (kbp), per technology |  |  |  |  |  |
| --- | --- | --- | --- | --- | --- | --- |
|  | PacBio CLR |  | PacBio HiFi |  | Oxford Nanopore |  |
|  | Mean | Stdev | Mean | Stdev | Mean | Stdev |
| **Iteration 1**<br>(technology-specific values) | 15.7 | 14.4 | 20.7 | 2.5 | 12.1 | 17.1 |
| **Iteration 2** | 25 | 22.5 | 25 | 3 | 25 | 35 |
| **Iteration 3**<br>(imitate ultra-long reads) | 35 | 31.5 | 35 | 4.2 | 35 | 49 |
| **Iteration 4** | 75 | 67.5 | 75 | 9 | 75 | 105 |

Consequently, we ran twelve simulations for each reference genome. As described above, we used our own models for each technology, and passed them to the simulator as the --error_model and --qscore_model. The read identities per technology were set to the values included in Supplementary table S2. Across all simulations, we chose a coverage depth of 30x. Canu’s documentation [18] specifies a minimum coverage of 20 - 25x for HiFi data, and 20x for other types of data, while Flye’s guidelines [19] indicate a minimum coverage of 30x. As there is no minimum recommended coverage indicated for the other assemblers we used in our benchmark, we simulated reads following the stricter guideline among these two, that is, 30x coverage.

A summary of the Badread commands used in our simulation can be found in Supplementary Table S3. Note that, in the case of simulated HiFi reads, we additionally lowered the rates of glitches, random, junk, and chimeric reads to reflect the higher accuracy of this technology. We set the percentage of chimeras to 0.04, as estimated by [20].

#### Real reads

In support of our evaluation on simulated reads, we also performed a benchmark on real-read assemblies from Oxford Nanopore and PacBio reads sequenced from the reference genomes. These reads were sampled to approximately 30x coverage, to ensure a fair comparison with our simulated-read assemblies. The data sources for all real sets are included in Supplementary Table S4.

### Assemblies

Five long-read de novo assemblers are included in this benchmark: Canu v2.2 [18], Flye v2.9 [19], Redbean (also known as Wtdbg2) v2.5 [21], Raven v1.7.0 [22], and Miniasm v0.3_r179 [23].

The assemblies were performed with default values for most parameters. Canu and Wtdbg2 require the estimated genome size as a parameter, and we set the following values: *S. cerevisiae* = 12 Mbp, *P. falciparum* = 23 Mbp, *A. thaliana* = 135 Mbp, *D. melanogaster* = 139 Mbp, *C. elegans* = 103 Mbp, and *T. rubripes* = 384 Mbp. All commands used in the assembly pipelines are available in Supplementary Table S6. We note that further polishing of assemblies using high-fidelity short reads, although common in practice [24–26], is omitted in this study, as the focus is exclusively on assembler performance on long-read data and not polishing tools.

We added a long-read polishing step for Miniasm and Wtdbg2, as their assembly pipelines do not include long-read based polishing. Following Raven’s default pipeline, which performs two rounds of Racon polishing [27], we used two rounds of Racon polishing on Wtdbg2 and Miniasm. We note that for Miniasm, we used Minipolish [7], which simplifies Racon polishing by applying it in two iterations on the GFA (Graphical Fragment Assembly) files produced by the assembler. For both Miniasm and Wtdbg2, the alignments required for polishing were generated with Minimap v2.24.

### Evaluation

We evaluated the assemblies in three different categories of metrics. The COMPASS analysis compares the assemblies with their corresponding reference genome and provides insight into their similarities. The assembly statistics provide some basic knowledge about the contiguity and misassemblies. Finally, the BUSCO assessment investigates the presence of essential genes in the assemblies. These three categories of metrics, next to each other, can provide a complete overview of the assembly’s quality.

### COMPASS analysis

For each assembly, we ran the COMPASS script to measure the coverage, validity, multiplicity and parsimony, to assess the quality of the assemblies, as defined in Assemblathon 2 [13]. These metrics describe several characteristics that were deemed important for comparing *de novo* assembly tools, and were computed using three types of data: (1) the reference sequence, (2) the assembled scaffolds, and (3) the alignments (sequences from the assembled scaffolds that were aligned to the reference sequences). Definitions and formulas for the metrics are reported in Supplementary Table S5.

#### Assembly statistics and misassembly events

We use QUAST v5.0.2 [28] is used to measure the NG50 [12] (Earl *et al*., 2011) of an assembly and the number of misassemblies. QUAST identifies misassemblies based on the definition outlined by [29]. The total number of misassemblies is the sum of all relocations, inversions, and translocations. Considering two adjacent flanking sequences, if they both align to the same chromosome, but 1 kbp away from each other, or overlapping for more than 1 kbp, this is counted as a relocation. If these flanking sequences, aligned to the same chromosome, are on opposite strands, the misassembly is considered an inversion. Lastly, translocations describe events in which two flanking sequences align to different chromosomes.

#### BUSCO assessment

BUSCO v5.4.2 assessment [30,31] is performed to evaluate the completeness of the essential genes in the assemblies. This quantifies the number of single-copy, duplicated, fragmented and missing orthologs in an assembled genome. From the number of orthologs specific to each dataset, BUSCO identifies how many orthologs are present in the assembly (either as single-copy or duplicated), how many are fragmented, and how many are missing. We ran these evaluations with a different OrthoDB lineage dataset for each genome: *S. cerevisiae* - saccharomycetes, *P. falciparum* - plasmodium, *A. thaliana* - brassicales, *D. melanogaster* - diptera, *C. elegans* - nematoda, and *T. rubripes* - actinopterygii.

## Results and discussion

### Overview of the benchmarking pipeline

Figure 1 shows an overview of the benchmarking pipeline. We begin with the selection of six representative eukaryotes from the interactive Tree of Life [11]: *S. cerevisiae, P. falciparum, A. thaliana, D. melanogaster, C. elegans*, and *T. rubripes*. We also use three read sets from the latest human assembly project [16] to generate Badread error and Qscore models [15] for PacBio Continuous Long Reads (CLRs), PacBio High Fidelity reads, and Oxford Nanopore reads (see Supplementary Table S2). The reference sequences and models become input to the Badread simulation stage. For each genome, we simulate reads with four different read length distributions and three sequencing technologies (see Table 1), amounting to a total of 12 simulated read sets per reference genome. These reads, as well as 13 real read sets, are assembled with five assembly tools: Canu, Flye, Miniasm, Raven, and Wtdbg2.

Next, the resulting assemblies are evaluated using COMPASS, QUAST, and BUSCO, and based on the reported metrics we distinguish six main evaluation categories: sequence identity, repeat collapse, rate of valid sequences, contiguity, misassembly count, and gene identification. The selected COMPASS metrics are the coverage, multiplicity, and validity of an assembly, which provide insight on sequence identity, repeat collapse, and the rate of valid sequences, respectively. In this regard, an ideal assembly has coverage, multiplicity and validity close to 1. This suggests that a large fraction of the reference genome is assembled, repeats are generally collapsed instead of replicated, and most sequences in the assembly are validated by the reference. Among others, QUAST reports the number of misassemblies and the NG50 of an assembly. A high NG50 value is associated with high contiguity. In order to assess contiguity across genomes of different sizes, we report the ratio between the assembly’s NG50 and the N50 of the references. Lastly, gene identification is quantified in terms of the percentage of complete BUSCOs in an assembly.

### The search for an optimal assembler is influenced by read sequencing technology, genome complexity, and research goal

To select an assembler that is most versatile across eukaryotic taxa, we simulate PacBio Continuous Long Reads (CLRs), PacBio High Fidelity (HiFi) reads, and Oxford Nanopore reads from the genomes of six model eukaryotes, assemble these reads, and evaluate the assemblers in the six main categories mentioned in the previous section. The results for each evaluation category are normalized in the range given by the worst and best values encountered in the evaluation of all assemblies of reads with default length. This highlights differences between assemblers, as well as between genomes and sequencing technologies.

The results of the benchmark on the read sets with default lengths, namely those belonging to the first iteration (see Table 1), are illustrated in Figure 2. A full report of the evaluation metrics in this figure is included in the Supplementary Tables S7 – S24, under “Iteration 1”. We note that no assembler unanimously ranks first in all categories, across different sequencing technologies and eukaryotic genomes, although our findings highlight some of their strengths and thus their potential for various research aims. The runtime and memory usage of the assembly tools on all of the simulated datasets are reported in Supplementary Tables S25 – S30, since this can also be a deciding factor next to the quality of the assembly for the researchers to choose the suitable assembler for their purpose. We note that all assemblies were run on our local High Performance Computing Cluster, and the runtime and RAM usage may have been affected by the heterogeneity of the shared computing environment in which the assembly jobs executed.

**Figure 2:**
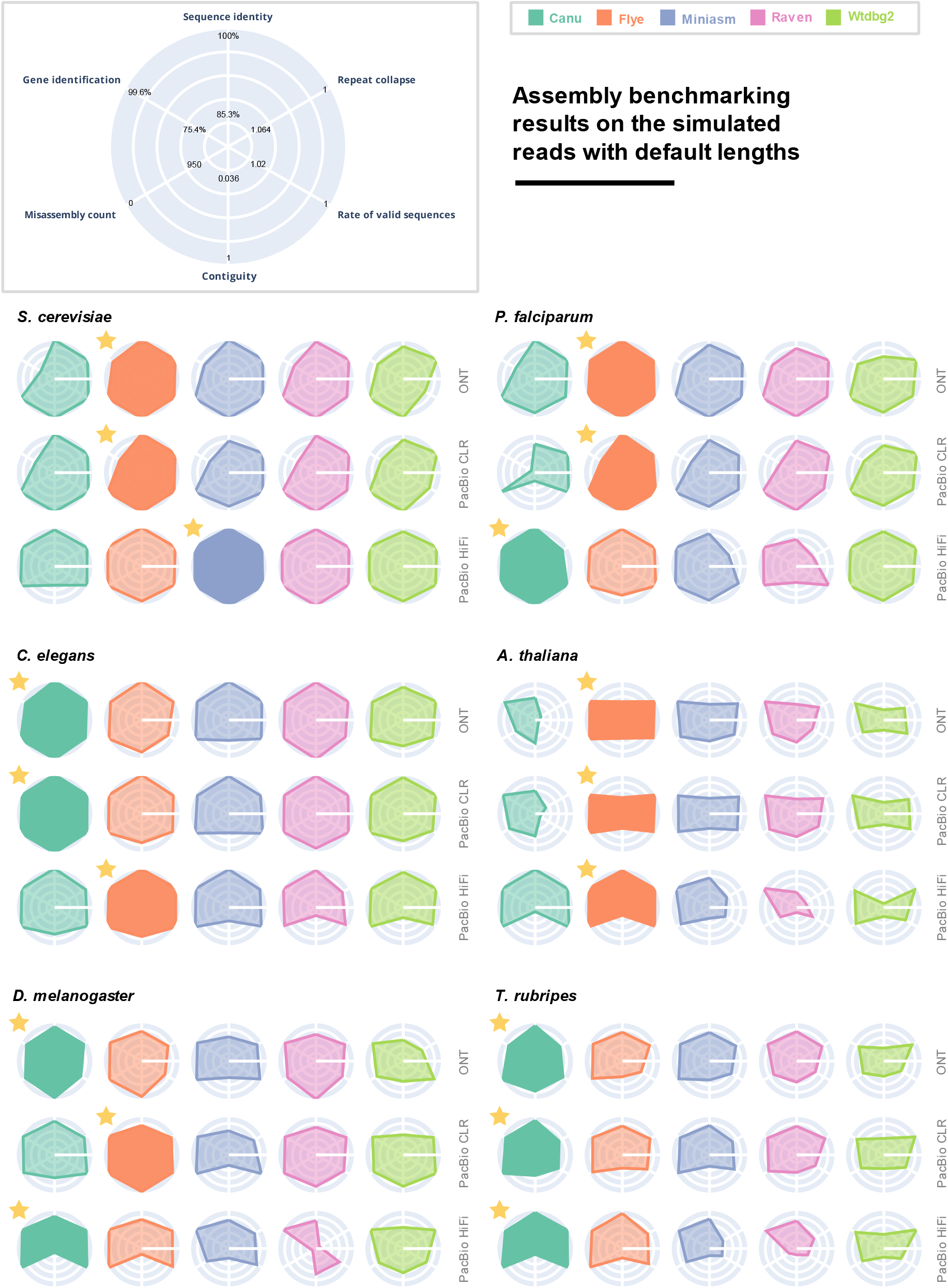
The performance of the five assemblers on the read sets with default read lengths, from iteration 1 (see Table 1), generated from six eukaryotic genomes. Six evaluation categories are reported for each assembler, and the results are normalized among all assemblies included in the figure. Ranges for each metric are reported as the best and worst values computed for these assemblies. The best performing assembler is highlighted for each read set, and marked with a star.

Miniasm, Raven and Wtdbg2 are all well-rounded choices for the simpler *S. cerevisiae, P. falciparum* and *C. elegans* genomes, with a balanced trade-off between assembly quality and computational resources. For PacBio HiFi reads, Raven is generally qualitatively outperformed by other assemblers like Canu, Flye, and Miniasm, likely as a consequence of the fact that its pipeline is not customized for all long-read sequencing technology. Nonetheless, if computational resources are a concern, Raven is a more suitable choice, since Miniasm and Wtdbg2 do not scale well for larger genomes.

We can single out Flye as the most robust assembler across all six organisms, although for larger genomes such as *T. rubripes*, Canu is a better tool. Both produce assemblies with high sequence identity and validity, as well as good gene prediction, but Flye assemblies generally rank first when we compute the average score across all six metrics. For Canu, we notice more variation in assembly quality across different genomes, particularly for *P. falciparum* and *A. thaliana*, while Flye maintains more consistent results. Nonetheless, on the *T. rubripes* genome, Canu assemblies have higher sequence identity and contiguity, as well as more accurate gene identification.

### Evaluation of real-read assemblies supports our rankings on simulated-read assemblies

To determine assembler performance on real reads and validate the rankings of the simulated-read assemblies, we assemble several real read sets from the six reference eukaryotes (Supplementary Table S4). The evaluation results on the real-read assemblies, summarized in Figure 3, indicate that assemblers which perform well on simulated reads perform similarly well in assembling the sets of real reads. The full report of metrics on the real read assemblies is included in Supplementary Table S31. We conclude that, overall, the assembler rankings remain consistent. This illustrates that benchmarking using simulated data is similar to real read sets. For reference-based metrics, we used the reference genomes given in Supplementary Table S1.

**Figure 3:**
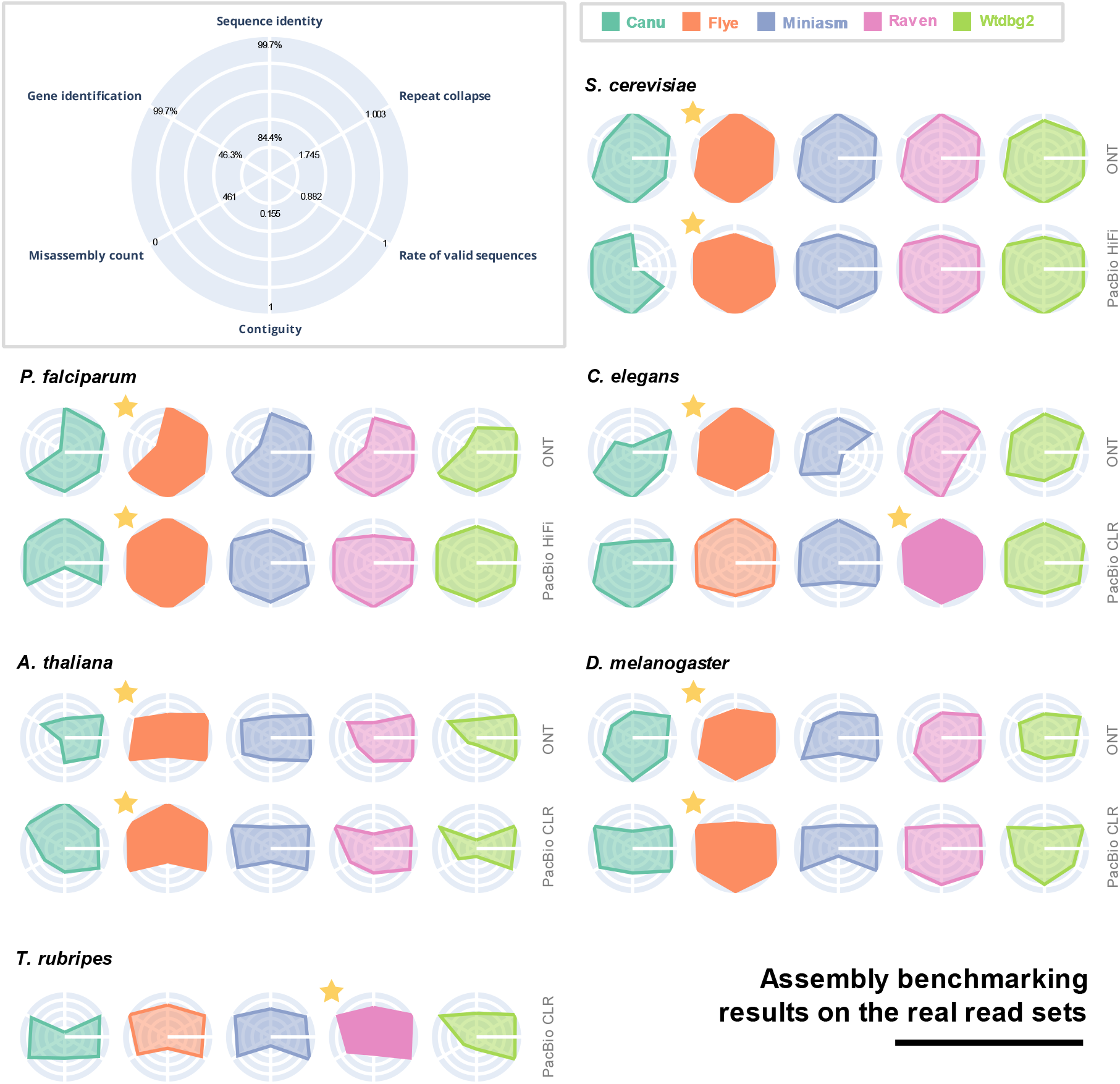
The performance of the five assemblers on the real reads (see Supplementary Table S4), sequenced from six eukaryotic genomes. As in Figure 2, six evaluation categories are reported for each assembler, and the results are normalized among all assemblies included in the figure. Ranges for each metric are reported as the best and worst values computed for these assemblies. The best performing assembler is highlighted for each read set, and marked with a star.

Notably, reference-based metrics in the evaluation of real-read assemblies rely on comparisons with an assembly, and not the genome from which the reads were initially sequenced. In contrast to the evaluation of simulated-read assemblies, the existence of a ground truth reference is not available in this case, but reference-based metrics are included for the sake of consistency with the simulated-read evaluation.

In the evaluation of real-read assemblies, Flye ranks first for nearly all datasets, with the exception of the *T. rubripes* and *C. elegans* PacBio reads, for which Raven performs better overall. However, even in *C. elegans*, Flye performance is close to the best values in all metrics other than contiguity. As expected, overall assembler performance decreases for reference-based metrics like sequence identity, repeat collapse and validity, but surprisingly the misassembly count is considerably lower.

### Longer reads lead to more contiguous assemblies of large genomes, but do not always improve assembly quality

To investigate the effect of increased read length on assembly quality, we use Badread to simulate Oxford Nanopore, as well as PacBio CLR and HiFi reads with different read length distributions (Table 1) from the genomes of *S. cerevisiae, P. falciparum, C. elegans, A. thaliana, D. melanogaster*, and *T. rubripes*. We assemble these reads with five state-of-the-art long-read assemblers, and evaluate assembly quality based on six evaluation categories (see Overview of the benchmarking pipeline). It is worth mentioning that Canu iteration 4 assemblies (the longest reads) of *A. thaliana* and *T. rubripes* did not finish within reasonable time and are excluded from the evaluation.

Figure 4 shows a summary of the assemblers’ performance on all simulated read sets, highlighting changes in performance for each read length distribution. All six evaluation metrics are normalized given the maximum and minimum metric values per genome, per sequencing technology, and combined to obtain an average score. We then average these three scores again and report a score between 1 and 10 for each assembler, per read length distribution. The results on all computed metrics are fully described in Supplementary Tables S7 – S24.

**Figure 4:**
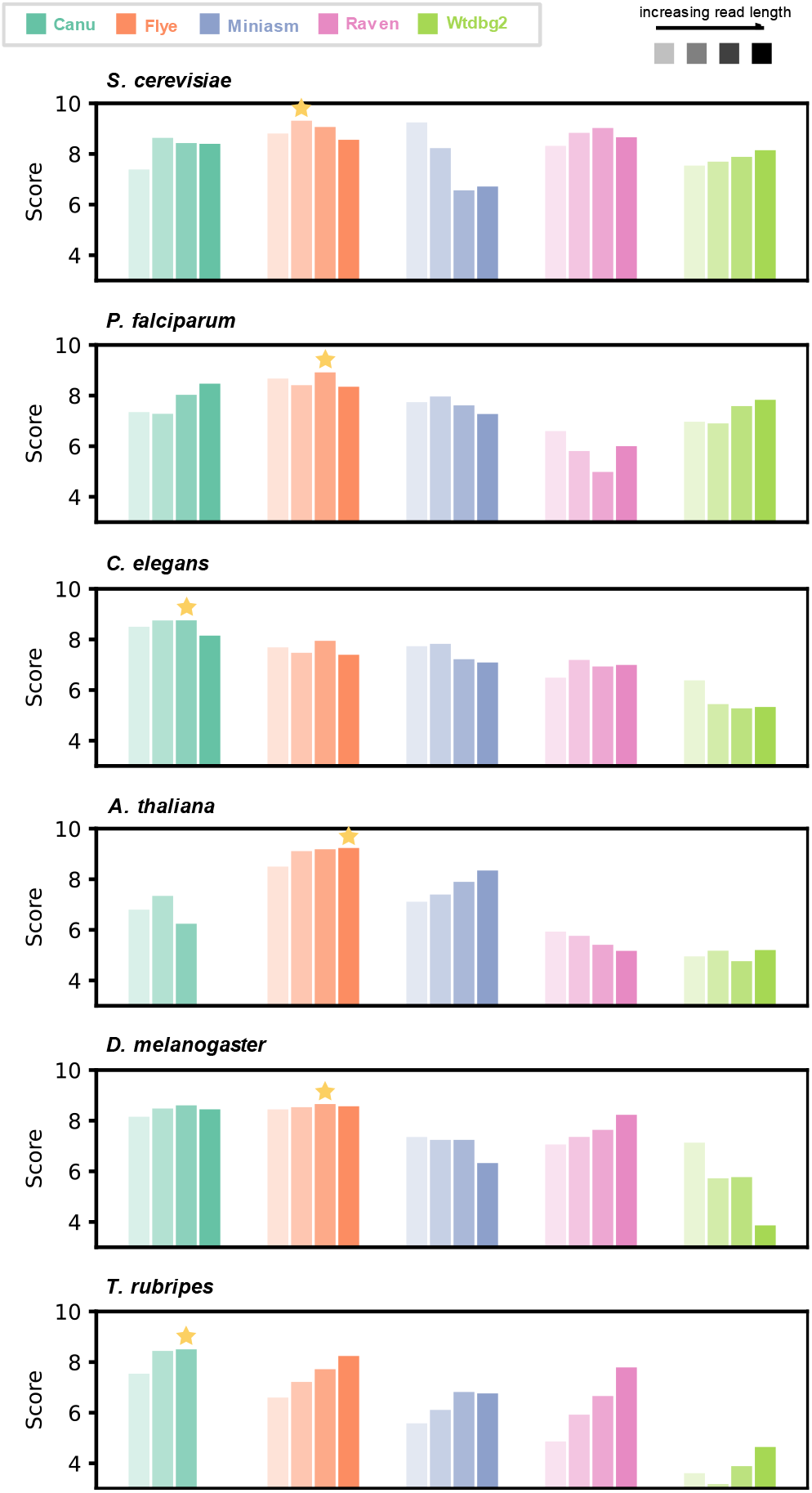
The performance of the five assemblers on all simulated read sets, with four different read length distributions (as previously described in Table 1). A score of 1 - 10 is reported for each assembler. The results are normalized for each genome, per sequencing technology. An average score for each read length distribution is first computed per technology (ONT, PacBio CLR, PacBio HiFi), and then these three scores are averaged to obtain an overall score per read length distribution.

The results imply that there is a correlation between the size and complexity of the reference genome and the extent of the improvement in assembly quality that can be achieved by increasing the length of the reads. While we observe no trend in assembly quality improvement on the assemblies of smaller genomes, the results on the *T. rubripes* assemblies are more conclusively in favour of the longer reads. For instance, on the shorter and simpler *S. cerevisiae* and *P. falciparum* genomes, identification of repetitive and complex regions is not aided by increased read length, likely as these regions are already spanned by the reads with default lengths. However, the benchmark results suggest that more complex and repetitive regions within the *A. thaliana, D. melanogaster* and, most notably, *T. rubripes* genomes are better captured by longer reads.

As recorded in Supplementary Table S22 – S24, for larger genomes, longer reads generally lead to significantly higher assembly contiguity and a lower misassembly count. The latter implies that the resulting assemblies are more faithful to the references, although this is not necessarily supported by other metrics. We cannot report any compelling improvements in sequence identity, multiplicity, validity, and gene identification.

## Conclusion

In fulfilment of the first objective of this study, we conclude that Flye is the highest performing assembler when considering the overview of all evaluation categories in this benchmark, which include the sequence identity, repeat collapse, rate of valid sequences, contiguity, misassembly count, and gene identification. Rankings are mostly consistent for all three sequencing platforms included in the study: PacBio CLR, PacBio HiFi and ONT. However, no assembler ranks first in all evaluation categories, suggesting that the choice of assembler is often a trade-off between certain advantages and disadvantages. Therefore, we have corroborated the conclusion of Wick and Holt [7], who benchmarked long-read assemblers on prokaryotes, for eukaryotic organisms, and recommend that these benchmarking parameters are considered in relation to the desired outcome of an assembly experiment.

Additionally, the tests performed on real reads validate our rankings of simulated-read assemblies. Flye, the assembler that scored consistently well in most evaluation categories for assemblies of simulated reads, also ranks first when evaluated on several sets of real reads sequenced on long-read platforms.

Regarding our second objective, which addressed the effect of increased read length on assembly quality, the benchmarking of assemblers on read sets with different read length distributions suggests that longer reads have the potential to improve assembly quality. However, this depends on the size and complexity of the genome that is being reconstructed. We found that improvements in contiguity were most significant among all metrics, as also supported by the conclusion of [8], who showed that using third generation sequencing considerably improves contiguity in assembling a plant genome (*M. jansenii*). However, we did not find significant improvements in other aspects of assembly quality, such as sequence identity or gene identification.

## Data availability

All accessions to the reference genomes used in this study are included in Supplementary Table S1. The read sets that were used for the creation of error and QScore models for the simulator are included in Supplementary Table S2. These models are available at https://github.com/AbeelLab/long-read-assembly-benchmark. The accessions for the real reads we assembled are included in Supplementary Table S4. All other data is reproducible as per the commands in Supplementary Tables S3 and S6.

## Code availability

Our evaluations were produced with QUAST v5.0.2 [28], BUSCO v5.4.2 [30, 31], and COMPASS [13]. We also provide the scripts we used on https://github.com/AbeelLab/long-read-assembly-benchmark.

