## Supplementary Table for "When do longer reads matter? A benchmark of long read de novo assembly tools for eukaryotic genomes"

### Supplementary Materials

**Supplementary Table S1:** Assembly accession numbers for all six reference genomes used in the experiments.

| **Organism** | **Reference assembly** |
|---|---|
| *S. cerevisiae* S288C | NCBI assembly R64, RefSeq accession GCF_000146045.2 |
| *A. thaliana* Ecotype Col-0 | NCBI assembly ASM2091176v1, GenBank accession GCA_020911765.1 (Hou *et al.*, 2022) |
| *C. elegans* VC2010 | ENA project accession PRJEB28388 (Yoshimura *et al.*, 2019) |
| *D. melanogaster* ISO-1 | NCBI RefSeq accession GCF_000001215.4 |
| *T. rubripes* | NCBI assembly fTakRub1.3, GenBank accession GCA_901000725.3 |
| *P. falciparum* (isolate: 3D7) | NCBI assembly GCA_000002765, GenBank accession GCA_000002765.3 |

**Supplementary Table S2:** Long read sets from the human genome used to generate Badread error and QScore models. Where needed, we downsampled reads to 3 Gbp, which meets the simulator's requirements for at least 1 Gbp of real sequence data. Read identities were calculated as described by (Wick, 2019), who used the definition of BLAST identity. The sequence data was aligned to reference GCF_009914755.1 (Nurk *et al.*, 2022), with Minimap v2.24 (Li, 2016).

| **Technology** | **Sequence data source** | **Read identities (%) (mean, max, stdev)** | **Notes** |
|---|---|---|---|
| PacBio Continuous Long Reads (CLRs) | HG002 extracted DNA: https://downloads.pacbcloud.com/public/dataset/SV-HG002-CLR/ | 89.349, 94.668, 2.678 | The read file was split into sub-files containing approximately 3 Gbp of data, using Fastaq. Alignment and training were performed using the first split. |
| PacBio Circular Consensus Sequencing (CCS) HiFi reads | Accession SRR11292120, from BioProject PRJNA530776 (Nurk *et al.*, 2022) | 99.748, 100.0, 0.291 | |
| Oxford Nanopore reads | https://s3.amazonaws.com/nanopore-human-wgs/chm13/nanopore/rel3/rel3.fastq.gz<br>Read IDs: https://s3.amazonaws.com/nanopore-human-wgs/chm13/nanopore/rel3/ids/ucd.ids.gz | 91.333, 98.783, 2.899 | The read file was split into sub-files containing approximately 3 Gbp of data, using Fastaq. Alignment and training were performed using the first split. |

**Supplementary Table S3:** Badread parameters used in the simulation of long reads, for each technology. In total, we simulated 84 read sets, accounting for 7 genomes (Supplementary Table S1), 3 sequencing technologies, and 4 read length distributions per technology (Table 1). Aside from read length, these parameters were kept consistent for each technology across all simulations. All other parameters not included in this table were kept as the simulator's defaults.

| | **Technology** | | |
|---|---|---|---|
| **Parameter** | **PacBio CLR** | **Oxford nanopore** | **PacBio HiFi** |
| **--error_model** | pacbio_human2019 (see Supplementary Table S2) | ont_human2019 (see Supplementary Table S2) | pacbio_hifi_human2022 (see Supplementary Table S2) |
| **--qscore_model** | pacbio_human2019 (see Supplementary Table S2) | ont_human2019 (see Supplementary Table S2) | pacbio_hifi_human2022 (see Supplementary Table S2) |
| **--seed** | 10 | | |
| **--identity** | 89.4,94.7,2.7 | 91.3,98.8,2.9 | 99.7,100.0,0.3 |
| **--length** | Varies per iteration (see Table 1) | | |
| **--quantity** | 30x | | |
| **--junk_reads** | default | | 0.01 |
| **--random_reads** | default | | 0.01 |
| **--chimeras** | default | | 0.04 |
| **--glitches** | default | | 100000,25,25 |

**Supplementary Table S4:** Accession for the sequencing data used in our benchmark of real-read assemblies. To match our simulated reads, we have further downsampled these read sets to 30x coverage.

| | Accessions | |
|---|---|---|
| **Organism** | **PacBio** | **Oxford Nanopore** |
| *S. cerevisiae* S288C | SRR18210286 | SRR17374240 |
| *A. thaliana* Ecotype Col-0 | SRR14728885 | SRR15720446 |
| *C. elegans* VC2010 | SRR7594465 | SRR7594463 |
| *D. melanogaster* ISO-1 | SRR11906525 | SRR6702603, SRR6821890 |
| *T. rubripes* | ERR3261643, ERR3253114, ERR3253113, ERR3253112 | - |
| *P. falciparum* (isolate: 3D7) | SRR13050273 | ERR10113962 |

**Supplementary Table S5:** Definitions and formulas for the COMPASS metrics defined in Assemblathon 2 (Bradnam *et al.*, 2013). We define $C, V, M, P$ as the coverage, validity, multiplicity, and parsimony of an assembly, respectively. We also denote $L_{CI}$ as the total length of the coverage islands, $L_A$ as the total length of the alignments between the reference and the assembly, $L_R$ as the total length of the reference, and $L_S$ as the total length of the assembly (sum of the scaffold lengths).

| **COMPASS metric** | **Definition** | **Formula** |
|---|---|---|
| Coverage | Coverage is a measure of the fraction of the reference genome that is present in an assembly. It is determined as the ratio between the summed length of the coverage islands and the summed length of the reference sequences. Coverage values range from 0 to 1, and a higher coverage is preferred. | $C = \frac{L_{CI}}{L_R}$ |
| Validity | Validity is the ratio between the summed length of the alignments and the assembled scaffolds, measuring how much of the assembly is aligned to the reference. By definition, a higher validity implies better assembly quality. | $V = \frac{L_A}{L_S}$ |

| | | |
|---|---|---|
| | However, validity values higher than 1 are encountered when the number of aligned bases is higher than the number of bases in the assembly, suggesting that some of the alignments overlap, implying that some alignments are duplicated. In this evaluation, we considered that a good validity score has a value close to 1. | |
| Multiplicity | Multiplicity is defined by the ratio of the summed length of the alignments and the summed length of the coverage islands. This metric gives insight on whether the assembler collapsed or replicated repeats within the genome. In the evaluation, multiplicity values close to 1 are considered better. | $M = \frac{L_A}{L_{CI}}$ |
| Parsimony | Parsimony is the ratio between multiplicity and validity. Low parsimony values are preferred, because this is interpreted as the assembly cost. The minimum parsimony value is 1, which means there is a one-to-one mapping from all of the assembly fragments to all of the sequences in the reference genome. | $P = \frac{M}{V}$ |

**Supplementary Table S6:** Assembly commands for all five assemblers. The `$genome_size` in the assembly commands below was set as follows: *S. cerevisiae* = 12 Mbp, *P. falciparum* = 23 Mbp, *A. thaliana* = 130 Mbp, *D. melanogaster* = 139 Mbp, *C. elegans* = 103 Mbp, and *T. rubripes* = 384 Mbp. The `$threads` parameter was set to 8 for *S. cerevisiae* and *P. falciparum*, 16 for *A. thaliana*, *C. elegans*, and *D. melanogaster*, and 20 for *T. rubripes*.

| **Assembler** | **Command** |
|---|---|
| Canu | `canu genomeSize=$genome_size -pacbio/-nanopore/-pacbio-hifi $reads MaxThreads=$threads useGrid=false -p $prefix -d $directory` |
| Flye | `flye --pacbio-hifi/--pacbio-raw/--nano-raw $reads --threads $threads --out-dir $directory` |
| Miniasm | `minimap2 -x ava-ont/ava-pb -t $threads $reads $reads \| gzip -1 > overlap.paf.gz`<br>`miniasm -f $reads overlap.paf.gz > unitigs.gfa`<br>`minipolish -t $threads $reads unitigs.gfa > assembly.gfa` |

| | |
|---|---|
| Raven | raven -t $threads $reads > assembly.fasta |
| Wtdbg2 | wtdbg2 -x ont/sq/ccs -g $genome_size -i $reads -t $threads -fo dbg<br>wtpoa-cns -t $threads -i dbg.ctg.lay.gz -fo dbg.raw.fa<br>minimap2 -t $threads -x map-ont/map-pb/map-hifi dbg.raw.fa $reads \| gzip > alignment1.paf.gz<br>racon -t $threads $reads alignment1.paf.gz dbg.raw.fa > polished1.fasta<br>minimap2 -t $threads -x map-ont/map-pb/map-hifi polished1.fasta $reads \| gzip > alignment2.paf.gz<br>racon -t $threads $reads alignment2.paf.gz polished1.fasta > assembly.fasta |

Supplementary Table S7: Evaluation results for the *S. cerevisiae* Oxford Nanopore simulated read assemblies.

***S. cerevisiae*: Oxford Nanopore reads**

| | Iteration 1 | Iteration 2 | Iteration 3 | Iteration 4 | |
|---|---|---|---|---|---|
| **Canu** | 0.999 | 0.993 | 0.999 | 0.93 | **Sequence identity** |
| **Flye** | 0.999 | 0.999 | 0.998 | 0.998 | |
| **Miniasm** | 0.999 | 0.931 | 0.843 | 0.674 | |
| **Raven** | 0.992 | 0.972 | 0.916 | 0.877 | |
| **Wtdbg2** | 0.972 | 0.965 | 0.966 | 0.954 | |
| **Canu** | 1.002 | 1 | 1.002 | 1 | **Repeat collapse** |
| **Flye** | 1.001 | 1.001 | 1.003 | 1.004 | |
| **Miniasm** | 1 | 1 | 1 | 1 | |
| **Raven** | 1.002 | 1.002 | 1 | 1.001 | |
| **Wtdbg2** | 1 | 1.001 | 1.008 | 1 | |
| **Canu** | 1 | 1.001 | 1.001 | 1.001 | **Rate of valid sequences** |
| **Flye** | 1.001 | 1.001 | 1.002 | 1.001 | |
| **Miniasm** | 1 | 1 | 1 | 1.002 | |
| **Raven** | 1.002 | 1 | 1.002 | 1.001 | |
| **Wtdbg2** | 0.992 | 0.999 | 0.988 | 0.999 | |
| **Canu** | 0.999 | 0.998 | 0.999 | 0.998 | **Contiguity (NG50 / N50)** |
| **Flye** | 1 | 1 | 1 | 1 | |
| **Miniasm** | 1 | 0.879 | 0.9 | 0.721 | |
| **Raven** | 0.999 | 0.999 | 0.999 | 0.999 | |
| **Wtdbg2** | 0.985 | 0.985 | 0.985 | 0.984 | |
| **Canu** | 0 | 0 | 0 | 0 | **Misassembly count** |
| **Flye** | 0 | 1 | 1 | 3 | |
| **Miniasm** | 0 | 3 | 0 | 2 | |
| **Raven** | 1 | 0 | 0 | 1 | |

| | | | | | |
|---|---|---|---|---|---|
| Wtdbg2 | 1 | 2 | 3 | 6 | |
| Canu | 83.5 | 84.1 | 84 | 79.1 | Gene identification (% complete BUSCOs) |
| Flye | 95.7 | 95.2 | 95.2 | 96.1 | |
| Miniasm | 92.1 | 87.9 | 80.2 | 64.6 | |
| Raven | 90.2 | 88.6 | 84.3 | 82.4 | |
| Wtdbg2 | 92.8 | 92.4 | 93 | 92.1 | |

Supplementary Table S8: Evaluation results for the S. cerevisiae PacBio CLR simulated read assemblies.

**S. cerevisiae: PacBio CLR reads**

| | Iteration 1 | Iteration 2 | Iteration 3 | Iteration 4 | |
|---|---|---|---|---|---|
| Canu | 0.998 | 0.998 | 0.998 | 0.993 | Sequence identity |
| Flye | 0.998 | 0.999 | 0.999 | 0.995 | |
| Miniasm | 0.969 | 0.883 | 0.835 | 0.683 | |
| Raven | 0.991 | 0.992 | 0.99 | 0.889 | |
| Wtdbg2 | 0.974 | 0.959 | 0.963 | 0.972 | |
| Canu | 1.002 | 1 | 1.002 | 1 | Repeat collapse |
| Flye | 1.001 | 1.002 | 1.004 | 1 | |
| Miniasm | 1 | 1.008 | 1.066 | 1 | |
| Raven | 1.002 | 1 | 1 | 1.001 | |
| Wtdbg2 | 1 | 1 | 1 | 1.002 | |
| Canu | 1 | 1.001 | 1.002 | 1.001 | Rate of valid sequences |
| Flye | 1.001 | 1.002 | 1.001 | 1 | |
| Miniasm | 1 | 0.997 | 0.999 | 1.002 | |
| Raven | 1.002 | 1 | 1 | 1.001 | |
| Wtdbg2 | 0.995 | 0.984 | 0.996 | 0.994 | |
| Canu | 0.999 | 0.999 | 0.999 | 0.999 | Contiguity (NG50 / N50) |
| Flye | 0.999 | 0.999 | 0.999 | 1.003 | |
| Miniasm | 0.848 | 0.806 | 0.634 | 0.807 | |
| Raven | 0.997 | 0.999 | 0.999 | 0.999 | |
| Wtdbg2 | 0.985 | 0.983 | 0.985 | 0.983 | |
| Canu | 0 | 0 | 0 | 0 | Misassembly count |
| Flye | 0 | 1 | 1 | 0 | |
| Miniasm | 0 | 0 | 2 | 0 | |
| Raven | 0 | 0 | 0 | 0 | |
| Wtdbg2 | 0 | 1 | 1 | 3 | |
| Canu | 87.5 | 87.1 | 88.9 | 88.5 | Gene identificati on (% [illegible] |
| Flye | 90 | 89.5 | 90.7 | 90.2 | |
| Miniasm | 87.8 | 81.4 | 76.9 | 64.5 | |

| | | | | |
|---|---|---|---|---|
| **Raven** | 87.1 | 87.5 | 88.7 | 81.3 |
| **Wtdbg2** | 89.9 | 89.4 | 89.8 | 89.4 |

Supplementary Table S9: Evaluation results for the S. cerevisiae PacBio HiFi simulated read assemblies.

**S. cerevisiae: PacBio Hifi reads**

| | **Iteration 1** | **Iteration 2** | **Iteration 3** | **Iteration 4** | |
|---|---|---|---|---|---|
| **Canu** | 0.989 | 0.987 | 0.985 | 0.976 | **Sequence identity** |
| **Flye** | 0.998 | 0.996 | 0.995 | 0.99 | |
| **Miniasm** | 0.996 | 0.993 | 0.994 | 0.991 | |
| **Raven** | 0.991 | 0.992 | 0.988 | 0.985 | |
| **Wtdbg2** | 0.982 | 0.983 | 0.985 | 0.988 | |
| **Canu** | 1.003 | 1.001 | 1.001 | 1 | **Repeat collapse** |
| **Flye** | 1.001 | 1.001 | 1 | 1 | |
| **Miniasm** | 1.002 | 1 | 1.001 | 1 | |
| **Raven** | 1.002 | 1.001 | 1.001 | 1 | |
| **Wtdbg2** | 1.001 | 1.001 | 1 | 1 | |
| **Canu** | 1.001 | 1 | 1 | 1 | **Rate of valid sequences** |
| **Flye** | 1.001 | 1 | 1 | 1.001 | |
| **Miniasm** | 1 | 1 | 1.001 | 1 | |
| **Raven** | 1.001 | 1.001 | 1 | 1 | |
| **Wtdbg2** | 1.001 | 1 | 1 | 1 | |
| **Canu** | 0.443 | 0.668 | 0.688 | 0.896 | **Contiguity (NG50 / N50)** |
| **Flye** | 0.88 | 1 | 0.873 | 1 | |
| **Miniasm** | 1 | 1.001 | 1 | 1 | |
| **Raven** | 1 | 1 | 1 | 0.993 | |
| **Wtdbg2** | 0.879 | 0.879 | 0.995 | 0.996 | |
| **Canu** | 0 | 0 | 0 | 0 | **Misassembly count** |
| **Flye** | 2 | 0 | 0 | 0 | |
| **Miniasm** | 0 | 1 | 1 | 0 | |
| **Raven** | 1 | 0 | 1 | 0 | |
| **Wtdbg2** | 4 | 4 | 0 | 1 | |
| **Canu** | 98.6 | 99 | 98.9 | 98.6 | **Gene identification (% complete BUSCOs)** |
| **Flye** | 99.6 | 99.6 | 99.3 | 99.2 | |
| **Miniasm** | 99.6 | 99.5 | 99.6 | 99.5 | |
| **Raven** | 99.6 | 99.6 | 99.6 | 99.5 | |
| **Wtdbg2** | 99.5 | 99.6 | 99.5 | 99.6 | |

Supplementary Table S10: Evaluation results for the P. falciparum Oxford Nanopore simulated read assemblies.

**P. falciparum: Oxford Nanopore reads**

| Metric | Assembler | Iteration 1 | Iteration 2 | Iteration 3 | Iteration 4 |
|---|---|---|---|---|---|
| Sequence identity | Canu | 0.998 | 0.997 | 0.999 | 0.995 |
| | Flye | 0.996 | 0.994 | 0.998 | 0.998 |
| | Miniasm | 0.979 | 0.979 | 0.96 | 0.952 |
| | Raven | 0.964 | 0.975 | 0.98 | 0.958 |
| | Wtdbg2 | 0.93 | 0.922 | 0.949 | 0.948 |
| Repeat collapse | Canu | 1.003 | 1.001 | 1.015 | 1.001 |
| | Flye | 1.005 | 1.008 | 1.006 | 1.004 |
| | Miniasm | 1.001 | 1.001 | 1 | 1.001 |
| | Raven | 1.003 | 1.005 | 1.007 | 1.004 |
| | Wtdbg2 | 1.002 | 1.004 | 1.003 | 1 |
| Rate of valid sequences | Canu | 0.998 | 0.996 | 1 | 1.002 |
| | Flye | 1.001 | 1.002 | 1.001 | 1.001 |
| | Miniasm | 1.001 | 1.001 | 1.001 | 1.001 |
| | Raven | 0.998 | 0.996 | 0.996 | 0.996 |
| | Wtdbg2 | 0.997 | 0.991 | 0.994 | 0.989 |
| Contiguity (NG50 / N50) | Canu | 0.854 | 0.854 | 0.998 | 0.997 |
| | Flye | 1 | 0.884 | 1 | 0.951 |
| | Miniasm | 0.978 | 0.981 | 1 | 0.982 |
| | Raven | 0.956 | 0.976 | 0.982 | 0.992 |
| | Wtdbg2 | 0.841 | 0.904 | 0.882 | 0.917 |
| Misassembly count | Canu | 0 | 0 | 2 | 0 |
| | Flye | 4 | 3 | 3 | 4 |
| | Miniasm | 0 | 0 | 0 | 1 |
| | Raven | 8 | 13 | 14 | 7 |
| | Wtdbg2 | 6 | 6 | 2 | 2 |
| Gene identification (% complete BUSCOs) | Canu | 87.6 | 88.6 | 89.4 | 89.2 |
| | Flye | 97.8 | 97.7 | 98.1 | 98 |
| | Miniasm | 92.5 | 92.4 | 90.4 | 90.7 |
| | Raven | 91.6 | 91.4 | 90.9 | 89.9 |
| | Wtdbg2 | 92.9 | 92.3 | 93.7 | 93.4 |

Supplementary Table S11: Evaluation results for the P. falciparum PacBio CLR simulated read assemblies.

**P. falciparum: PacBio CLR reads**

| Metric | Assembler | Iteration 1 | Iteration 2 | Iteration 3 | Iteration 4 |
|---|---|---|---|---|---|
| Sequence identity | Canu | 0.956 | 0.959 | 0.975 | 0.974 |
| | Flye | 0.999 | 0.994 | 0.996 | 0.996 |
| | Miniasm | 0.973 | 0.937 | 0.934 | 0.836 |
| | Raven | 0.97 | 0.975 | 0.98 | 0.983 |
| | Wtdbg2 | 0.95 | 0.954 | 0.961 | 0.96 |
| Repeat collapse | Canu | 1.002 | 1.003 | 1.001 | 1.001 |
| | Flye | 1.006 | 1.007 | 1.004 | 1.005 |
| | Miniasm | 1.009 | 1.012 | 1.011 | 1.012 |
| | Raven | 1.002 | 1.004 | 1.006 | 1.005 |
| | Wtdbg2 | 1.001 | 1 | 1 | 1 |
| Rate of valid sequences | Canu | 1 | 1.002 | 1.004 | 1.004 |
| | Flye | 1 | 0.999 | 1.003 | 1.003 |
| | Miniasm | 0.997 | 1 | 1 | 0.999 |
| | Raven | 0.996 | 0.992 | 0.993 | 0.973 |
| | Wtdbg2 | 0.997 | 0.993 | 0.992 | 0.99 |
| Contiguity (NG50 / N50) | Canu | 0.156 | 0.282 | 0.432 | 0.54 |
| | Flye | 1 | 0.993 | 0.947 | 1 |
| | Miniasm | 0.853 | 0.818 | 0.856 | 0.854 |
| | Raven | 0.955 | 0.993 | 0.981 | 0.997 |
| | Wtdbg2 | 0.826 | 0.868 | 0.863 | 0.893 |
| Misassembly count | Canu | 0 | 0 | 0 | 0 |
| | Flye | 2 | 3 | 2 | 2 |
| | Miniasm | 0 | 0 | 0 | 1 |
| | Raven | 4 | 9 | 7 | 4 |
| | Wtdbg2 | 5 | 3 | 1 | 0 |
| Gene identification (% complete BUSCOs) | Canu | 75.4 | 77.2 | 79.6 | 79 |
| | Flye | 88.9 | 88.1 | 88.1 | 88.4 |
| | Miniasm | 86.8 | 84.9 | 83.3 | 76.7 |
| | Raven | 85.3 | 85.2 | 85.9 | 85.3 |
| | Wtdbg2 | 88.1 | 87.5 | 87.8 | 87.4 |

Supplementary Table S12: Evaluation results for the P. falciparum PacBio HiFi simulated read assemblies.

**P. falciparum: PacBio HiFi reads**

| Metric | Assembler | Iteration 1 | Iteration 2 | Iteration 3 | Iteration 4 |
|---|---|---|---|---|---|
| Sequence identity | Canu | 0.999 | 1 | 0.998 | 0.997 |
| | Flye | 0.992 | 0.997 | 0.996 | 0.982 |
| | Miniasm | 0.973 | 0.979 | 0.984 | 0.987 |
| | Raven | 0.949 | 0.941 | 0.942 | 0.959 |
| | Wtdbg2 | 0.982 | 0.977 | 0.98 | 0.989 |
| Repeat collapse | Canu | 1.013 | 1.019 | 1.01 | 1.012 |
| | Flye | 1.003 | 1.002 | 1.003 | 1.001 |
| | Miniasm | 1.03 | 1.021 | 1.014 | 1.001 |
| | Raven | 1.036 | 1.055 | 1.123 | 1.08 |
| | Wtdbg2 | 1.003 | 1.002 | 1.002 | 1.001 |
| Rate of valid sequences | Canu | 1.001 | 1.001 | 1 | 1 |
| | Flye | 1 | 1 | 1.001 | 1 |
| | Miniasm | 1.003 | 1.001 | 1.003 | 1.001 |
| | Raven | 0.999 | 0.998 | 0.997 | 0.997 |
| | Wtdbg2 | 1.003 | 1.002 | 1.002 | 1.001 |
| Contiguity (NG50 / N50) | Canu | 0.996 | 0.914 | 0.914 | 1 |
| | Flye | 0.716 | 0.914 | 0.924 | 0.856 |
| | Miniasm | 0.854 | 0.874 | 0.893 | 0.984 |
| | Raven | 0.368 | 0.309 | 0.188 | 0.609 |
| | Wtdbg2 | 0.864 | 0.878 | 0.883 | 0.931 |
| Misassembly count | Canu | 1 | 1 | 0 | 0 |
| | Flye | 1 | 1 | 2 | 2 |
| | Miniasm | 11 | 5 | 2 | 0 |
| | Raven | 47 | 29 | 53 | 24 |
| | Wtdbg2 | 2 | 5 | 0 | 1 |
| Gene identification (% complete BUSCOs) | Canu | 98.6 | 98.4 | 98.7 | 98.6 |
| | Flye | 98 | 98.5 | 98.6 | 97.6 |
| | Miniasm | 98.7 | 98.7 | 98.7 | 98.7 |
| | Raven | 98.6 | 98.6 | 98.4 | 98.7 |
| | Wtdbg2 | 98.7 | 98.6 | 98.6 | 98.6 |

Supplementary Table S13: Evaluation results for the C. elegans Oxford Nanopore simulated read assemblies.

**C. elegans: Oxford Nanopore reads**

| Metric | Assembler | Iteration 1 | Iteration 2 | Iteration 3 | Iteration 4 |
|---|---|---|---|---|---|
| Sequence identity | Canu | 1 | 1 | 0.999 | 0.998 |
| | Flye | 0.998 | 0.998 | 0.998 | 0.998 |
| | Miniasm | 0.996 | 0.991 | 0.989 | 0.99 |
| | Raven | 0.994 | 0.997 | 0.996 | 0.996 |
| | Wtdbg2 | 0.974 | 0.984 | 0.955 | 0.951 |
| Repeat collapse | Canu | 1.004 | 1.004 | 1.003 | 1.002 |
| | Flye | 1.005 | 1.007 | 1.005 | 1.01 |
| | Miniasm | 1.002 | 1.002 | 1.003 | 1.002 |
| | Raven | 1.004 | 1.003 | 1.007 | 1.003 |
| | Wtdbg2 | 1.003 | 1.002 | 1.002 | 1.005 |
| Rate of valid sequences | Canu | 1.002 | 1.001 | 0.997 | 1.004 |
| | Flye | 1.005 | 1.005 | 1.003 | 1.004 |
| | Miniasm | 1.001 | 1.001 | 1.002 | 1.002 |
| | Raven | 1.003 | 1.002 | 1.003 | 0.994 |
| | Wtdbg2 | 0.997 | 0.995 | 0.996 | 0.995 |
| Contiguity (NG50 / N50) | Canu | 0.997 | 0.994 | 0.994 | 0.993 |
| | Flye | 0.807 | 1.002 | 0.999 | 0.996 |
| | Miniasm | 0.487 | 0.858 | 0.962 | 0.996 |
| | Raven | 0.997 | 0.998 | 0.873 | 0.872 |
| | Wtdbg2 | 0.638 | 0.365 | 0.629 | 0.761 |
| Misassembly count | Canu | 3 | 6 | 5 | 8 |
| | Flye | 37 | 36 | 29 | 49 |
| | Miniasm | 22 | 33 | 53 | 34 |
| | Raven | 56 | 43 | 36 | 55 |
| | Wtdbg2 | 38 | 27 | 39 | 72 |
| Gene identification (% complete BUSCOs) | Canu | 95.7 | 96 | 95.9 | 95 |
| | Flye | 97.4 | 97.5 | 97.8 | 97.5 |
| | Miniasm | 97.9 | 97.3 | 97.4 | 96.5 |
| | Raven | 97.4 | 97.5 | 97 | 97.2 |
| | Wtdbg2 | 97.2 | 97.6 | 95.7 | 93.9 |

Supplementary Table S14: Evaluation results for the C. elegans PacBio CLR simulated read assemblies.

**C. elegans: PacBio CLR reads**

| Metric | Assembler | Iteration 1 | Iteration 2 | Iteration 3 | Iteration 4 |
|---|---|---|---|---|---|
| Sequence identity | Canu | 1 | 1 | 1 | 0.999 |
| | Flye | 0.998 | 0.998 | 0.998 | 0.998 |
| | Miniasm | 0.993 | 0.993 | 0.988 | 0.975 |
| | Raven | 0.994 | 0.995 | 0.997 | 0.997 |
| | Wtdbg2 | 0.988 | 0.992 | 0.989 | 0.985 |
| Repeat collapse | Canu | 1.003 | 1.001 | 1.003 | 1.003 |
| | Flye | 1.005 | 1.005 | 1.005 | 1.009 |
| | Miniasm | 1.005 | 1.008 | 1.016 | 1.009 |
| | Raven | 1.004 | 1.003 | 1.004 | 1.003 |
| | Wtdbg2 | 1.003 | 1.003 | 1.003 | 1.004 |
| Rate of valid sequences | Canu | 1.001 | 0.999 | 0.999 | 1.001 |
| | Flye | 0.998 | 0.997 | 0.998 | 1.002 |
| | Miniasm | 1 | 1.001 | 1 | 1 |
| | Raven | 1.001 | 1.001 | 1.002 | 1.001 |
| | Wtdbg2 | 0.997 | 0.996 | 0.995 | 0.995 |
| Contiguity (NG50 / N50) | Canu | 1 | 1 | 1 | 1 |
| | Flye | 0.722 | 0.863 | 0.874 | 1.017 |
| | Miniasm | 0.451 | 0.715 | 0.338 | 0.583 |
| | Raven | 0.872 | 0.999 | 1 | 1 |
| | Wtdbg2 | 0.672 | 0.749 | 0.788 | 0.997 |
| Misassembly count | Canu | 10 | 10 | 4 | 1 |
| | Flye | 21 | 33 | 40 | 30 |
| | Miniasm | 7 | 6 | 3 | 10 |
| | Raven | 53 | 40 | 39 | 55 |
| | Wtdbg2 | 22 | 31 | 34 | 50 |
| Gene identification (% complete BUSCOs) | Canu | 98.5 | 98.4 | 98.5 | 98.1 |
| | Flye | 98.7 | 98.7 | 98.7 | 98.7 |
| | Miniasm | 97.7 | 98 | 97.9 | 95.6 |
| | Raven | 97.9 | 98.3 | 98.2 | 98.1 |
| | Wtdbg2 | 98.2 | 98.3 | 98.1 | 97.1 |

Supplementary Table S15: Evaluation results for the C. elegans PacBio HiFi simulated read assemblies.

**C. elegans: PacBio HiFi reads**

| Metric | Assembler | Iteration 1 | Iteration 2 | Iteration 3 | Iteration 4 |
|---|---|---|---|---|---|
| Sequence identity | Canu | 0.999 | 0.999 | 0.999 | 1 |
| | Flye | 0.996 | 0.997 | 0.997 | 0.997 |
| | Miniasm | 0.995 | 0.995 | 0.995 | 0.995 |
| | Raven | 0.992 | 0.993 | 0.995 | 0.993 |
| | Wtdbg2 | 0.984 | 0.964 | 0.973 | 0.996 |
| Repeat collapse | Canu | 1.006 | 1.003 | 1.003 | 1.105 |
| | Flye | 1 | 1.001 | 1.001 | 1.002 |
| | Miniasm | 1.006 | 1.005 | 1.003 | 1.002 |
| | Raven | 1.016 | 1.016 | 1.012 | 1.009 |
| | Wtdbg2 | 1.002 | 1.003 | 1.002 | 1.003 |
| Rate of valid sequences | Canu | 1.001 | 1.001 | 1.001 | 1.002 |
| | Flye | 1 | 1.001 | 1.001 | 1.001 |
| | Miniasm | 1.001 | 1.002 | 1.001 | 1.001 |
| | Raven | 1.003 | 1.004 | 1.003 | 1.001 |
| | Wtdbg2 | 1.003 | 1.004 | 1.003 | 1.002 |
| Contiguity (NG50 / N50) | Canu | 0.659 | 0.82 | 1.863 | 3.216 |
| | Flye | 0.723 | 0.722 | 0.82 | 0.819 |
| | Miniasm | 0.295 | 0.722 | 0.819 | 0.722 |
| | Raven | 0.15 | 0.279 | 0.211 | 0.386 |
| | Wtdbg2 | 0.206 | 0.219 | 0.44 | 1 |
| Misassembly count | Canu | 14 | 11 | 10 | 12 |
| | Flye | 1 | 7 | 3 | 5 |
| | Miniasm | 12 | 11 | 10 | 0 |
| | Raven | 76 | 52 | 33 | 27 |
| | Wtdbg2 | 16 | 17 | 10 | 13 |
| Gene identification (% complete BUSCOs) | Canu | 98.7 | 98.8 | 98.7 | 98.8 |
| | Flye | 98.8 | 98.8 | 98.7 | 98.7 |
| | Miniasm | 98.8 | 98.8 | 98.8 | 98.7 |
| | Raven | 98.5 | 98.5 | 98.6 | 98.4 |
| | Wtdbg2 | 98.3 | 96.8 | 96.8 | 98.8 |

Supplementary Table S16: Evaluation results for the A. thaliana Oxford Nanopore simulated read assemblies.

**A. thaliana: Oxford Nanopore reads**

| | Iteration 1 | Iteration 2 | Iteration 3 | Iteration 4 | |
|---|---|---|---|---|---|
| **Canu** | 0.929 | 0.934 | 0.937 | | Sequence identity |
| **Flye** | 0.913 | 0.92 | 0.925 | 0.936 | |
| **Miniasm** | 0.902 | 0.906 | 0.909 | 0.914 | |
| **Raven** | 0.902 | 0.909 | 0.91 | 0.915 | |
| **Wtdbg2** | 0.879 | 0.885 | 0.859 | 0.871 | |
| **Canu** | 1.059 | 1.055 | 1.108 | | Repeat collapse |
| **Flye** | 1.004 | 1.003 | 1.004 | 1.005 | |
| **Miniasm** | 1.012 | 1.013 | 1.012 | 1.007 | |
| **Raven** | 1.025 | 1.035 | 1.036 | 1.043 | |
| **Wtdbg2** | 1.028 | 1.068 | 1.086 | 1.105 | |
| **Canu** | 1.02 | 1.017 | 1.028 | | Rate of valid sequences |
| **Flye** | 1.002 | 1.002 | 1.003 | 1.002 | |
| **Miniasm** | 1.006 | 1.007 | 1.006 | 1.002 | |
| **Raven** | 1.012 | 1.016 | 1.014 | 1.013 | |
| **Wtdbg2** | 1.007 | 1.008 | 1.008 | 1.004 | |
| **Canu** | 0.563 | 0.571 | 0.58 | | Contiguity (NG50 / N50) |
| **Flye** | 0.43 | 0.56 | 0.574 | 0.573 | |
| **Miniasm** | 0.505 | 0.558 | 0.522 | 0.525 | |
| **Raven** | 0.535 | 0.524 | 0.535 | 0.573 | |
| **Wtdbg2** | 0.205 | 0.206 | 0.114 | 0.218 | |
| **Canu** | 461 | 423 | 871 | | Misassembly count |
| **Flye** | 30 | 9 | 7 | 9 | |
| **Miniasm** | 159 | 155 | 111 | 6 | |
| **Raven** | 312 | 459 | 467 | 596 | |
| **Wtdbg2** | 405 | 388 | 424 | 335 | |
| **Canu** | 96.7 | 96.6 | 96.5 | | Gene identification (% complete BUSCOs) |
| **Flye** | 98.4 | 98.6 | 98.5 | 98.5 | |
| **Miniasm** | 98 | 98 | 98 | 98 | |
| **Raven** | 97.7 | 97.8 | 97.5 | 97.5 | |
| **Wtdbg2** | 95.1 | 95 | 92.4 | 93.1 | |

Supplementary Table S17: Evaluation results for the A. thaliana PacBio CLR simulated read assemblies.

**A. thaliana: PacBio CLR reads**

| | Iteration 1 | Iteration 2 | Iteration 3 | Iteration 4 | |
|---|---|---|---|---|---|
| Canu | 0.932 | 0.931 | 0.936 | 0.935 | Sequence identity |
| Flye | 0.904 | 0.909 | 0.91 | 0.919 | |
| Miniasm | 0.901 | 0.898 | 0.896 | 0.891 | |
| Raven | 0.898 | 0.902 | 0.904 | 0.906 | |
| Wtdbg2 | 0.884 | 0.885 | 0.878 | 0.889 | |
| Canu | 1.049 | 1.03 | 1.047 | 1.047 | Repeat collapse |
| Flye | 1.003 | 1.003 | 1.007 | 1.006 | |
| Miniasm | 1.01 | 1.008 | 1.011 | 1.014 | |
| Raven | 1.016 | 1.019 | 1.028 | 1.033 | |
| Wtdbg2 | 1.019 | 1.018 | 1.026 | 1.052 | |
| Canu | 1.019 | 1.012 | 1.017 | 1.015 | Rate of valid sequences |
| Flye | 0.998 | 0.999 | 1.002 | 1.004 | |
| Miniasm | 1.004 | 1.003 | 1.002 | 1.001 | |
| Raven | 1.008 | 1.01 | 1.011 | 1.013 | |
| Wtdbg2 | 1.005 | 1.001 | 1.002 | 0.999 | |
| Canu | 0.539 | 0.562 | 0.562 | 0.573 | Contiguity (NG50 / N50) |
| Flye | 0.329 | 0.536 | 0.562 | 0.56 | |
| Miniasm | 0.329 | 0.431 | 0.506 | 0.556 | |
| Raven | 0.557 | 0.587 | 0.558 | 0.588 | |
| Wtdbg2 | 0.225 | 0.175 | 0.224 | 0.486 | |
| Canu | 257 | 225 | 406 | 397 | Misassembly count |
| Flye | 19 | 23 | 19 | 50 | |
| Miniasm | 94 | 82 | 57 | 64 | |
| Raven | 215 | 232 | 287 | 374 | |
| Wtdbg2 | 285 | 259 | 237 | 231 | |
| Canu | 98.1 | 97.8 | 98.1 | 98 | Gene identification (% complete BUSCOs) |
| Flye | 98.5 | 98.8 | 98.7 | 98.6 | |
| Miniasm | 97.4 | 97.2 | 97.6 | 96.7 | |
| Raven | 97.4 | 97.3 | 97.4 | 97.1 | |
| Wtdbg2 | 97.4 | 96.4 | 96.2 | 97.1 | |

Supplementary Table S18: Evaluation results for the A. thaliana PacBio Hifi simulated read assemblies.

**A. thaliana: PacBio HiFi reads**

| Metric | Assembler | Iteration 1 | Iteration 2 | Iteration 3 | Iteration 4 |
|---|---|---|---|---|---|
| Sequence identity | **Canu** | 0.995 | 0.995 | 0.996 | 0.995 |
| Sequence identity | **Flye** | 0.991 | 0.992 | 0.992 | 0.982 |
| Sequence identity | **Miniasm** | 0.96 | 0.967 | 0.977 | 0.992 |
| Sequence identity | **Raven** | 0.9 | 0.9 | 0.91 | 0.951 |
| Sequence identity | **Wtdbg2** | 0.853 | 0.877 | 0.889 | 0.871 |
| Repeat collapse | **Canu** | 1.003 | 1.002 | 1.003 | 1.006 |
| Repeat collapse | **Flye** | 1.003 | 1.003 | 1.003 | 1.002 |
| Repeat collapse | **Miniasm** | 1.035 | 1.038 | 1.034 | 1.022 |
| Repeat collapse | **Raven** | 1.053 | 1.048 | 1.056 | 1.061 |
| Repeat collapse | **Wtdbg2** | 1.006 | 1.005 | 1.004 | 1.007 |
| Rate of valid sequences | **Canu** | 1 | 1 | 1.001 | 1 |
| Rate of valid sequences | **Flye** | 1.001 | 1.002 | 1.002 | 1.001 |
| Rate of valid sequences | **Miniasm** | 1.012 | 1.011 | 1.009 | 1.006 |
| Rate of valid sequences | **Raven** | 1.012 | 1.013 | 1.017 | 1.02 |
| Rate of valid sequences | **Wtdbg2** | 1.008 | 1.008 | 1.006 | 1.007 |
| Contiguity (NG50 / N50) | **Canu** | 0.045 | 0.057 | 0.096 | 0.272 |
| Contiguity (NG50 / N50) | **Flye** | 0.143 | 0.189 | 0.29 | 0.309 |
| Contiguity (NG50 / N50) | **Miniasm** | 0.224 | 0.288 | 0.433 | 0.558 |
| Contiguity (NG50 / N50) | **Raven** | 0.068 | 0.07 | 0.079 | 0.073 |
| Contiguity (NG50 / N50) | **Wtdbg2** | 0.174 | 0.235 | 0.337 | 0.512 |
| Misassembly count | **Canu** | 4 | 4 | 4 | 0 |
| Misassembly count | **Flye** | 7 | 11 | 9 | 1 |
| Misassembly count | **Miniasm** | 172 | 169 | 134 | 40 |
| Misassembly count | **Raven** | 548 | 501 | 592 | 520 |
| Misassembly count | **Wtdbg2** | 191 | 152 | 129 | 93 |
| Gene identification (% complete BUSCOs) | **Canu** | 98.7 | 98.7 | 99 | 98.9 |
| Gene identification (% complete BUSCOs) | **Flye** | 99.2 | 99.2 | 99.2 | 99 |
| Gene identification (% complete BUSCOs) | **Miniasm** | 99.2 | 99.3 | 99.3 | 99.2 |
| Gene identification (% complete BUSCOs) | **Raven** | 99.1 | 99.1 | 99.2 | 98.6 |
| Gene identification (% complete BUSCOs) | **Wtdbg2** | 95.6 | 97.8 | 98.3 | 96.2 |

Supplementary Table S19: Evaluation results for the D. melanogaster Oxford Nanopore simulated read assemblies.

**D. melanogaster: Oxford Nanopore reads**

| Metric | Assembler | Iteration 1 | Iteration 2 | Iteration 3 | Iteration 4 |
|---|---|---|---|---|---|
| Sequence identity | Canu | 0.98 | 0.98 | 0.981 | 0.982 |
| | Flye | 0.961 | 0.965 | 0.966 | 0.964 |
| | Miniasm | 0.938 | 0.935 | 0.922 | 0.868 |
| | Raven | 0.948 | 0.953 | 0.951 | 0.949 |
| | Wtdbg2 | 0.924 | 0.93 | 0.914 | 0.921 |
| Repeat collapse | Canu | 1.01 | 1.012 | 1.014 | 1.01 |
| | Flye | 1.016 | 1.011 | 1.013 | 1.012 |
| | Miniasm | 1.012 | 1.01 | 1.013 | 1.031 |
| | Raven | 1.011 | 1.009 | 1.01 | 1.01 |
| | Wtdbg2 | 1.032 | 1.119 | 1.148 | 1.189 |
| Rate of valid sequences | Canu | 1.005 | 1.003 | 0.998 | 0.999 |
| | Flye | 1.007 | 1.006 | 1.007 | 1.008 |
| | Miniasm | 1.002 | 1.003 | 1.001 | 1 |
| | Raven | 1.005 | 1.002 | 1.004 | 0.999 |
| | Wtdbg2 | 0.998 | 0.996 | 0.992 | 0.989 |
| Contiguity (NG50 / N50) | Canu | 0.929 | 0.999 | 0.968 | 0.999 |
| | Flye | 0.898 | 0.928 | 0.93 | 0.933 |
| | Miniasm | 0.379 | 0.645 | 0.497 | 0.34 |
| | Raven | 0.965 | 0.929 | 0.997 | 1.109 |
| | Wtdbg2 | 0.481 | 0.379 | 0.5 | 0.185 |
| Misassembly count | Canu | 148 | 131 | 145 | 168 |
| | Flye | 53 | 29 | 43 | 49 |
| | Miniasm | 52 | 41 | 27 | 34 |
| | Raven | 173 | 182 | 178 | 230 |
| | Wtdbg2 | 238 | 501 | 614 | 681 |
| Gene identification (% complete BUSCOs) | Canu | 95 | 95 | 94.3 | 93.5 |
| | Flye | 97.8 | 98 | 97.9 | 97.6 |
| | Miniasm | 97.6 | 97.8 | 97.5 | 94.7 |
| | Raven | 97.2 | 97.3 | 97.3 | 97.6 |
| | Wtdbg2 | 96.7 | 94.8 | 93.2 | 92.5 |

Supplementary Table S20: Evaluation results for the D. melanogaster PacBio CLR simulated read assemblies.

**D. melanogaster: PacBio CLR reads**

| Metric | Assembler | Iteration 1 | Iteration 2 | Iteration 3 | Iteration 4 |
|---|---|---|---|---|---|
| Sequence identity | Canu | 0.977 | 0.977 | 0.973 | 0.974 |
| | Flye | 0.959 | 0.963 | 0.964 | 0.966 |
| | Miniasm | 0.937 | 0.931 | 0.903 | 0.841 |
| | Raven | 0.952 | 0.954 | 0.953 | 0.954 |
| | Wtdbg2 | 0.92 | 0.937 | 0.93 | 0.921 |
| Repeat collapse | Canu | 1.009 | 1.008 | 1.008 | 1.007 |
| | Flye | 1.009 | 1.008 | 1.01 | 1.013 |
| | Miniasm | 1.019 | 1.037 | 1.033 | 1.045 |
| | Raven | 1.008 | 1.007 | 1.005 | 1.008 |
| | Wtdbg2 | 1.008 | 1.019 | 1.054 | 1.129 |
| Rate of valid sequences | Canu | 1.001 | 1.002 | 1.002 | 1.004 |
| | Flye | 0.998 | 0.997 | 1.002 | 1.004 |
| | Miniasm | 0.999 | 0.998 | 0.997 | 0.999 |
| | Raven | 1.004 | 1.004 | 1.003 | 0.998 |
| | Wtdbg2 | 0.999 | 0.999 | 0.992 | 0.984 |
| Contiguity (NG50 / N50) | Canu | 0.557 | 0.969 | 0.999 | 0.998 |
| | Flye | 0.934 | 0.9 | 0.901 | 0.928 |
| | Miniasm | 0.216 | 0.144 | 0.148 | 0.156 |
| | Raven | 0.793 | 0.918 | 1 | 0.97 |
| | Wtdbg2 | 0.789 | 0.851 | 0.796 | 0.751 |
| Misassembly count | Canu | 98 | 88 | 104 | 111 |
| | Flye | 36 | 18 | 36 | 40 |
| | Miniasm | 39 | 42 | 41 | 54 |
| | Raven | 69 | 55 | 33 | 53 |
| | Wtdbg2 | 143 | 134 | 206 | 375 |
| Gene identification (% complete BUSCOs) | Canu | 97.6 | 97.3 | 98 | 97.3 |
| | Flye | 98.4 | 98.1 | 98.5 | 98.4 |
| | Miniasm | 97.3 | 97 | 96.3 | 91.2 |
| | Raven | 97.2 | 96.5 | 97.3 | 97.2 |
| | Wtdbg2 | 96.9 | 97 | 96.1 | 94 |

Supplementary Table S21: Evaluation results for the D. melanogaster PacBio HiFi simulated read assemblies.

**D. melanogaster: PacBio HiFi reads**

| Metric | Assembler | Iteration 1 | Iteration 2 | Iteration 3 | Iteration 4 |
|---|---|---|---|---|---|
| Sequence identity | Canu | 0.971 | 0.97 | 0.971 | 0.973 |
| | Flye | 0.958 | 0.959 | 0.96 | 0.955 |
| | Miniasm | 0.955 | 0.953 | 0.953 | 0.948 |
| | Raven | 0.953 | 0.954 | 0.956 | 0.958 |
| | Wtdbg2 | 0.925 | 0.9 | 0.935 | 0.92 |
| Repeat collapse | Canu | 1.005 | 1.005 | 1.004 | 1.008 |
| | Flye | 1.007 | 1.007 | 1.005 | 1.004 |
| | Miniasm | 1.017 | 1.017 | 1.011 | 1.015 |
| | Raven | 1.064 | 1.06 | 1.053 | 1.038 |
| | Wtdbg2 | 1.004 | 1.004 | 1.003 | 1.006 |
| Rate of valid sequences | Canu | 1.001 | 1.001 | 1 | 1.001 |
| | Flye | 1.002 | 1.003 | 1.002 | 1.001 |
| | Miniasm | 1.004 | 1.005 | 1.003 | 1.002 |
| | Raven | 1.007 | 1.006 | 1.006 | 1.003 |
| | Wtdbg2 | 1.007 | 1.006 | 1.005 | 1.003 |
| Contiguity (NG50 / N50) | Canu | 0.037 | 0.053 | 0.071 | 0.17 |
| | Flye | 0.086 | 0.137 | 0.248 | 0.303 |
| | Miniasm | 0.19 | 0.22 | 0.261 | 0.611 |
| | Raven | 0.621 | 0.502 | 0.49 | 0.508 |
| | Wtdbg2 | 0.591 | 0.252 | 0.86 | 0.896 |
| Misassembly count | Canu | 29 | 38 | 29 | 19 |
| | Flye | 31 | 18 | 22 | 21 |
| | Miniasm | 377 | 295 | 286 | 224 |
| | Raven | 950 | 847 | 650 | 491 |
| | Wtdbg2 | 303 | 289 | 271 | 276 |
| Gene identification (% complete BUSCOs) | Canu | 98.2 | 98.1 | 98.4 | 98.4 |
| | Flye | 98.6 | 98.7 | 98.7 | 98.6 |
| | Miniasm | 98.8 | 98.7 | 98.8 | 98.8 |
| | Raven | 98.7 | 98.7 | 98.7 | 98.7 |
| | Wtdbg2 | 98.5 | 95.4 | 98.5 | 95.6 |

Supplementary Table S22: Evaluation results for the T. rubripes Oxford Nanopore simulated read assemblies.

**T. rubripes: ONT reads**

| Metric | Assembler | Iteration 1 | Iteration 2 | Iteration 3 | Iteration 4 |
|---|---|---|---|---|---|
| Sequence identity | **Canu** | 0.984 | 0.985 | 0.983 | |
| | **Flye** | 0.957 | 0.965 | 0.967 | 0.97 |
| | **Miniasm** | 0.956 | 0.948 | 0.944 | 0.914 |
| | **Raven** | 0.96 | 0.965 | 0.969 | 0.967 |
| | **Wtdbg2** | 0.891 | 0.892 | 0.9 | 0.915 |
| Repeat collapse | **Canu** | 1.019 | 1.01 | 1.013 | |
| | **Flye** | 1.014 | 1.013 | 1.013 | 1.018 |
| | **Miniasm** | 1.016 | 1.013 | 1.007 | 1.005 |
| | **Raven** | 1.018 | 1.017 | 1.018 | 1.017 |
| | **Wtdbg2** | 1.012 | 1.025 | 1.031 | 1.043 |
| Rate of valid sequences | **Canu** | 1.004 | 1.003 | 1.006 | |
| | **Flye** | 1.01 | 1.01 | 1.008 | 1.009 |
| | **Miniasm** | 1.008 | 1.005 | 1.003 | 1.004 |
| | **Raven** | 1.01 | 1.007 | 1.006 | 1.002 |
| | **Wtdbg2** | 1.011 | 1.009 | 1.008 | 1.002 |
| Contiguity (NG50 / N50) | **Canu** | 0.75 | 0.909 | 0.957 | |
| | **Flye** | 0.357 | 0.526 | 0.811 | 0.883 |
| | **Miniasm** | 0.445 | 0.674 | 0.766 | 0.826 |
| | **Raven** | 0.502 | 0.698 | 0.899 | 0.871 |
| | **Wtdbg2** | 0.344 | 0.401 | 0.367 | 0.368 |
| Misassembly count | **Canu** | 48 | 29 | 41 | |
| | **Flye** | 123 | 99 | 102 | 101 |
| | **Miniasm** | 102 | 38 | 30 | 44 |
| | **Raven** | 339 | 248 | 209 | 164 |
| | **Wtdbg2** | 412 | 377 | 382 | 381 |
| Gene identification (% complete BUSCOs) | **Canu** | 93.8 | 93.6 | 93 | |
| | **Flye** | 95.9 | 95.5 | 95.6 | 95.5 |
| | **Miniasm** | 95.8 | 95 | 94.5 | 92.4 |
| | **Raven** | 95.5 | 95.3 | 95.1 | 95.5 |
| | **Wtdbg2** | 92.3 | 91.7 | 92.6 | 92.9 |

Supplementary Table S23: Evaluation results for the T. rubripes PacBio CLR simulated read assemblies.

**T. rubripes: PacBio CLR reads**

| Metric | Assembler | Iteration 1 | Iteration 2 | Iteration 3 | Iteration 4 |
|---|---|---|---|---|---|
| Sequence identity | Canu | 0.98 | 0.979 | 0.979 | 0.971 |
| | Flye | 0.957 | 0.959 | 0.962 | 0.968 |
| | Miniasm | 0.959 | 0.957 | 0.955 | 0.931 |
| | Raven | 0.956 | 0.963 | 0.966 | 0.962 |
| | Wtdbg2 | 0.905 | 0.892 | 0.929 | 0.917 |
| Repeat collapse | Canu | 1.021 | 1.013 | 1.01 | 1.008 |
| | Flye | 1.013 | 1.009 | 1.01 | 1.011 |
| | Miniasm | 1.023 | 1.024 | 1.018 | 1.021 |
| | Raven | 1.012 | 1.012 | 1.011 | 1.012 |
| | Wtdbg2 | 1.006 | 1.011 | 1.015 | 1.028 |
| Rate of valid sequences | Canu | 1.007 | 1.005 | 1.003 | 1.003 |
| | Flye | 1.006 | 1.006 | 1.006 | 1.007 |
| | Miniasm | 1.006 | 1.006 | 1.003 | 1.002 |
| | Raven | 1.01 | 1.007 | 1.006 | 1.004 |
| | Wtdbg2 | 1.008 | 1.009 | 1.009 | 1.001 |
| Contiguity (NG50 / N50) | Canu | 0.492 | 0.683 | 0.808 | 0.81 |
| | Flye | 0.292 | 0.434 | 0.445 | 0.702 |
| | Miniasm | 0.242 | 0.409 | 0.49 | 0.618 |
| | Raven | 0.41 | 0.535 | 0.804 | 0.81 |
| | Wtdbg2 | 0.309 | 0.414 | 0.467 | 0.445 |
| Misassembly count | Canu | 57 | 28 | 24 | 16 |
| | Flye | 132 | 105 | 109 | 93 |
| | Miniasm | 93 | 89 | 52 | 45 |
| | Raven | 182 | 148 | 86 | 53 |
| | Wtdbg2 | 348 | 315 | 294 | 280 |
| Gene identification (% complete BUSCOs) | Canu | 96.1 | 96.3 | 96.1 | 95.2 |
| | Flye | 96.2 | 96.6 | 96.5 | 96.6 |
| | Miniasm | 96 | 95.5 | 95.1 | 93.7 |
| | Raven | 95.7 | 95.9 | 95.7 | 94.8 |
| | Wtdbg2 | 94.1 | 92.4 | 95.1 | 93.2 |

Supplementary Table S24: Evaluation results for the T. rubripes PacBio HiFi simulated read assemblies.

**T. rubripes: PacBio HiFi reads**

| | Iteration 1 | Iteration 2 | Iteration 3 | Iteration 4 | |
|---|---|---|---|---|---|
| **Canu** | 0.987 | 0.987 | 0.986 | 0.988 | **Sequence identity** |
| **Flye** | 0.983 | 0.983 | 0.984 | 0.984 | |
| **Miniasm** | 0.96 | 0.963 | 0.969 | 0.975 | |
| **Raven** | 0.952 | 0.955 | 0.961 | 0.968 | |
| **Wtdbg2** | 0.896 | 0.893 | 0.894 | 0.921 | |
| **Canu** | 1.004 | 1.003 | 1.003 | 1.008 | **Repeat collapse** |
| **Flye** | 1.018 | 1.015 | 1.011 | 1.003 | |
| **Miniasm** | 1.043 | 1.042 | 1.044 | 1.037 | |
| **Raven** | 1.035 | 1.034 | 1.037 | 1.025 | |
| **Wtdbg2** | 1.003 | 1.003 | 1.003 | 1.004 | |
| **Canu** | 1.001 | 1.001 | 1 | 1 | **Rate of valid sequences** |
| **Flye** | 1.005 | 1.004 | 1.003 | 1.002 | |
| **Miniasm** | 1.014 | 1.013 | 1.011 | 1.007 | |
| **Raven** | 1.015 | 1.012 | 1.013 | 1.006 | |
| **Wtdbg2** | 1.011 | 1.011 | 1.01 | 1.011 | |
| **Canu** | 0.087 | 0.128 | 0.154 | 0.429 | **Contiguity (NG50 / N50)** |
| **Flye** | 0.185 | 0.223 | 0.305 | 0.602 | |
| **Miniasm** | 0.123 | 0.152 | 0.202 | 0.409 | |
| **Raven** | 0.101 | 0.114 | 0.141 | 0.38 | |
| **Wtdbg2** | 0.164 | 0.216 | 0.302 | 0.641 | |
| **Canu** | 30 | 27 | 19 | 14 | **Misassembly count** |
| **Flye** | 59 | 51 | 36 | 21 | |
| **Miniasm** | 343 | 271 | 234 | 90 | |
| **Raven** | 811 | 731 | 551 | 247 | |
| **Wtdbg2** | 391 | 365 | 295 | 243 | |
| **Canu** | 96.4 | 96.5 | 96.5 | 96.7 | **Gene identification (% complete BUSCOs)** |
| **Flye** | 96.8 | 96.9 | 96.8 | 96.5 | |
| **Miniasm** | 96.8 | 96.8 | 96.8 | 96.9 | |
| **Raven** | 96.8 | 96.9 | 96.8 | 96.7 | |
| **Wtdbg2** | 95 | 94.1 | 93.7 | 94.8 | |
